# No *Wolbachia* protection against *Plasmodium falciparum* infection in the major malaria mosquito *Anopheles moucheti*

**DOI:** 10.1101/2023.05.28.542560

**Authors:** Théo Mouillaud, Audric Berger, Marie Buysse, Nil Rahola, Josquin Daron, Jean-Pierre Agbor, Sandrine N. Sango, Daniel E. Neafsey, Olivier Duron, Diego Ayala

## Abstract

Since the discovery of natural malaria vector populations infected by the endosymbiont bacterium *Wolbachia*, a renewed interest has arisen for using this bacterium as an alternative for malaria control. Among naturally infected mosquitoes, *Anopheles moucheti*, a major malaria mosquito in Central Africa, exhibits one of the highest prevalences of *Wolbachia* infection. To better understand whether this maternally inherited bacterium could be used for malaria control, we investigated *Wolbachia* influence in *An. moucheti* populations naturally infected by the malaria parasite *Plasmodium falciparum*. To this end, we collected mosquitoes in a village from Cameroon, Central Africa, where this mosquito is the main malaria vector. We found that the prevalence of *Wolbachia* bacterium was almost fixed in the studied mosquito population, and was higher than previously recorded. We also quantified *Wolbachia* in whole mosquitoes and dissected abdomens, confirming that the bacterium is also elsewhere than in the abdomen, but at lower density. Finally, we analyzed the impact of *Wolbachia* presence and density on *P. falciparum* infection. *Wolbachia* density was slightly higher in mosquitoes infected with the malaria parasite than in uninfected mosquitoes. However, we observed no correlation between the *P. falciparum* and *Wolbachia* densities. In conclusion, our study indicates that naturally occurring *Wolbachia* infection does not affect *P. falciparum* development within *An. moucheti* mosquitoes.

## Introduction

In the last decades, *Wolbachia* has emerged as a promising tool against vector-borne diseases in addition to the classical control measures, such as the use of insecticides (Bourtzis et al., 2014; Caragata, Dutra, Sucupira, Ferreira, & Moreira, 2021; Ferguson, 2018; Iturbe-Ormaetxe, Walker, & Neill, 2011; Ross, Turelli, & Hoffmann, 2019). This endosymbiont bacterium presents two major characteristics that can be exploited as an alternative vector control strategy. First, *Wolbachia* bacteria can induce cytoplasmic incompatibility in their hosts to enhance their maternal transmission. Specifically, the host sperm-egg compatibility is altered when infected males mate with uninfected females (Werren, Baldo, & Clark, 2008). Usually, this reproductive phenotype exhibits high prevalence in their hosts (Caragata et al., 2021; Ross et al., 2019). Therefore, releasing infected males in a non-infected population should drastically reduce the vector density, called population suppression. This approach has been successfully employed to control dengue transmission by the mosquito *Aedes aegypti* (Crawford et al., 2020; Utarini et al., 2021). Second, this endosymbiont bacterium can mediate and protect their hosts from infection by a pathogen. This protective phenotype has been observed for a large variety of pathogens and hosts (Braquart-Varnier et al., 2015; Cattel, Martinez, Jiggins, Mouton, & Gibert, 2016; Gupta et al., 2017; Martinez et al., 2014). Nevertheless, the underlying molecular mechanisms are still unknown. Some studies suggest a competition for the host cell resources that limits the parasite development (Paredes, Herren, Schüpfer, & Lemaitre, 2016). However, other studies hypothesized the pre-activation of the host innate immune system (Iturbe-Ormaetxe et al., 2011; Pan et al., 2018). This implies that *Wolbachia* infection stimulates the host active defenses, and consequently has a protective effect against pathogens. This phenotype has been observed mainly when *Wolbachia* is artificially transferred into a new host (Ross et al., 2019). In natural systems, the prevalence of an endosymbiont with a protective phenotype infection can occur at intermediate or low frequencies if the intensity of reproductive manipulation is low (Hilgenboecker, Hammerstein, Schlattmann, Telschow, & Werren, 2008; Ross et al., 2019). The two phenotypes are not exclusive, and they have been concomitantly observed upon the release of *Ae. aegypti* infected with the *Wolbachia* strain *w*Mel (Ross et al., 2022; Ross et al., 2019).

In *Anopheles* mosquitoes, the discovery of *Wolbachia* infection in natural conditions has been controversial. For decades, it was thought that *Anopheles* vectors were not infected by this bacterium due to the presence of *Asaia* as antagonist to *Wolbachia* (Grant L Hughes et al., 2014; Rossi et al., 2015). However, few years ago, *Wolbachia* was detected in *Anopheles gambiae*, the major malaria vector in Africa (Baldini et al., 2014). The low infection frequency (i.e. <20%) may explain the previous failures to detect these bacteria, but led to doubts about this finding (Chrostek & Gerth, 2018; Sawadogo et al., 2022). However, other authors confirmed the presence of *Wolbachia* in this and other *Anopheles* species (Ayala et al., 2019; Jeffries et al., 2018; Walker et al., 2021), and how *Wolbachia* interact with *Anopheles* microbiota (Straub et al., 2020). *Wolbachia* was already studied in *Anopheles* by transfer of an exogenous strain into *An. gambiae* and *Anopheles stephensi* (Bian et al., 2013; Grant L. Hughes, Koga, Xue, Fukatsu, & Rasgon, 2011; Kambris et al., 2010). Despite the promising effect on blocking malaria transmission, it was impossible to establish a self-sustainable infection by maternal inheritance (Grant L Hughes et al., 2014), limiting its applicability. After the discovery of naturally occurring infected *An. gambiae* and *Anopheles coluzzii* populations, some authors investigated *Wolbachia* role in *Plasmodium falciparum* infection and revealed a strong negative correlation between *Wolbachia* presence and *P. falciparum* infection (Gomes et al., 2017; Shaw et al., 2016). Although most *Anopheles* species exhibit a low *Wolbachia* infection rate, between 0% and 20% (Ayala et al., 2019), few species show higher rates, particularly *Anopheles moucheti* (50 to 70% in independent populations), in which vertical transmission was confirmed (Ayala et al., 2019; Walker et al., 2021). *Anopheles moucheti* sensu stricto (hereafter *An. moucheti*) is considered one of the major malaria vectors in Africa (C. Antonio-Nkondjio et al., 2006; Fontenille & Simard, 2004). It belongs to a group of 3 species, among which *An. moucheti* is the main malaria vector (Christophe Antonio-Nkondjio & Simard, 2013). This malaria mosquito is considered to be at the origin of human malaria from the transfer of the non-human malaria parasites from primates to humans (Paupy et al., 2013). Moreover, it is broadly present in the forested areas of Central Africa (Ayala et al., 2009), where it contributes significantly to malaria transmission. Therefore, the high *Wolbachia* prevalence in this mosquito species offers a compelling opportunity to study how this endosymbiont bacterium modulates *Plasmodium* infection in its host. Unfortunately, one of the major drawbacks to experimental approaches is the absence of established *An. moucheti* laboratory colonies, limiting current investigations to mosquitoes sampled from natural populations.

In this study, we investigated the effect of *Wolbachia* on *P. falciparum* infection in a natural *An. moucheti* population in Cameroon, Central Africa. We screened mosquitoes to detect the presence of *Wolbachia* and *P. falciparum* using a newly developed qPCR assay. We further quantified the density of both microorganisms in *An. moucheti* specimens. With this approach, we empirically assessed the *Wolbachia* effect on *P. falciparum* infection in this mosquito under natural conditions. Our results contribute to the study of *Wolbachia* in *Anopheles* and its potential value to the development of new strategies for malaria control.

## Material and Methods

### Sample collection, mosquito identification, and DNA extraction

Mosquitoes were collected in Ndangueng, Cameroon, in February 2020 and July 2021. Adult females were sampled using the human landing catch method (HLC), following the recommendations of the National Ethical Committee in Cameroon (CNERSH N° 2020/07/1259/CE/CNERSH/SP). Mosquitoes were morphologically identified on the basis of taxonomical identification keys (M. T. Gillies & Coetzee, 1987; Michael Thomas Gillies & De Meillon, 1968). Specimens belonging to the *An. moucheti* group were transported and kept alive for up to 10 days at the Malaria research Service of OCEAC, Yaoundé, Cameroon. For mosquitoes that died within the first 3 days after capture (gonotrophic cycle), abdomens were dissected and kept separately. All dead specimens were individually preserved in tubes containing a desiccant (silica gel, Sigma-Aldrich) and stored at -20°C for molecular studies. DNA was extracted using the CTAB method, as described in Morlais *et al* (Morlais, Poncon, Simard, Cohuet, & Fontenille, 2004). Briefly, samples were ground in 200µl of 2% CTAB solution (1M Tris HCl pH 8.0, 0.5M EDTA, 1.4 M NaCl, 2% cetyltrimethyl ammonium bromide) and incubated at 65°C for 5 min. Total DNA was extracted in chloroform, precipitated in isopropanol, washed in 70% ethanol before resuspension in Qiagen TE buffer and storage at -20°C. Then, a sample subset was analyzed to identify members of the *An. moucheti* group using the species-specific PCR assays developed by Kengne *et al*., (Kengne et al., 2007).

### Wolbachia and P. falciparum detection and quantification

The detection and quantification of *Wolbachia* and *P. falciparum* in *An. moucheti* mosquitoes were assessed using quantitative PCR assays (here after qPCR). To this end, a plasmid carrying a specific fragment of gene from both targeted organisms was designed. All primers were designed with Primer3plus (Untergasser et al., 2012), targeting mono-copy housekeeping genes to avoid over-estimations: *An. moucheti GPDG* gene (https://www.ebi.ac.uk/, assembly: GCA_943734755.1) (F: 5’-AAGTTGTTTCCGGACGTTTG-3’; R: 5’-CGTCGGATAGATTAATGGTG-3’), *Wolbachia coxA* gene (GenBank accession number: MK755519.1) (F: 5’-GGTGCTATAACTATGCTGCT-3’; R: 5’-TATGTAAACTTCTGGATGACC-3’), and *P. falciparum Cox1* gene (Genbank accession number: LR605957.1) (F: 5’-TTACATCAGGAATGTTATTGC -3’; R: 5’-ATATTGGATCTCCTGCAAAT -3’)(Boissiere et al., 2013). To construct the plasmid, the three amplicons (*An. moucheti*, 111bp; *Wolbachia*, 126bp; *P. falciparum*, 120 bp) were cloned together (total length of 357 bp) (Text S1) in the pEX-A128 vector. The primers, amplicons and plasmid were prepared by Eurofins.

The molecular quantification of *P. falciparum* and *Wolbachia* gene copies was carried out by qPCR. 1µl of total DNA was added to a mixture (final volume of 10µl) that contained 0.6 µl of each primer (see above) at 10µM, 0.5 µl of 2X Mix SensiFast Sybr NO-ROX Kit (Bioline), and 2.8µl of sterile molecular biology grade water (Hyclone Hypure, Cytiva). Each plasmid contains a gene copy for each organism, therefore, calculating the number of plasmid copies, we can estimate the gene copies for both *Wolbachia* and *P. falciparum* in each mosquito. To construct the standard curve, the plasmid was diluted seven times from 10^−2^ to 10^−7^ (7.45×10^7^ to 7.45×10^2^ plasmid copies/µl). For each PCR plate, one separate mixture for each of the three targeted genes per sample and per plasmid dilution was prepared. In total, 24 samples were tested in each 96 PCR plate. Cycling conditions included an initial denaturation step at 95°C for 10 min, followed by 40 cycles of denaturation (95°C for 10 s), annealing (57°C for 5 s), and elongation (72°C for 20 s), followed by a melting step (95°C for 2 min, 68°C for 2 min, and then up to 97°C, holding for 15 s once at 97°C) with a continuous fluorescence detection to construct the melting curves, and a final cooling cycle at 40°C for 10 s. All qPCR assays were carried out on a LightCycler 480 (Roche Diagnostics) using the LightCycler 480 software version 1.5.1.

### Statistical analysis and data visualization

Statistical analyses were carried out with R (R Core Team, 2021) and data modeling with the package “*tidyverse*” (Wickham et al., 2019). To estimate the repeatability of our quantitative measures, the mixed effects model framework implemented by the package “*rptR*”(Stoffel, Nakagawa, & Schielzeth, 2020) was used with the “species” variable (i.e. *P. falciparum* or *Wolbachia*) as random-effect predictors. Groups (i.e. infected vs non-infected mosquitoes) were compared with the Mann–Whitney test and the package “stats”. Correlation between ratios (i.e. *P. falciparum* and *Wolbachia*) were computed using the Pearson correlation coefficient (*r*) in R. Data, including all figures, were visualized using the package “*ggplot*” (Wickham, 2016) and associated packages, such as “*patchwork*” (Pedersen, 2020) and “*ggpubr*” (Kassambara & Kassambara, 2020).

## Results

### Prevalence of malaria parasites and Wolbachia in An. moucheti

In total, we collected 2042 mosquitoes belonging to the *An. moucheti* group (Table S1). Although, *An. moucheti* is considered the predominant species of the group in this region (Ayala et al., 2009), we molecularly analyzed a subset of 93 mosquitoes to determine the species (Kengne et al., 2007) and attest that they were all assigned to the *An. moucheti* species. Therefore, we considered that *An. moucheti* was the only species of the group present in our sampling. To analyze their role in malaria transmission, the prevalence of *P. falciparum* infected mosquitoes was determined through qPCR assays. We found that 64/2042 females (3%) were infected with *P. falciparum* (Fig 1A), a slightly higher rate than what previously reported (∼1.4%) in the same geographic zone (Christophe Antonio-Nkondjio et al., 2002). We then assessed *Wolbachia* presence in the infected mosquitoes and equal number of non-infected to *Plasmodium*. In total, we analyzed 130 specimens. We removed specimens showing elevated cycle threshold (*ct* > 26) to avoid any potential false positive amplification. Among the 113 specimens retained, 107 (95%) were infected by *Wolbachia* (Fig 1B). This rate was higher than what previously observed in Gabon (71%, (Ayala et al., 2019)) and Cameroon (56.6%, (Walker et al., 2021)). We used these 113 mosquitoes for the subsequent analysis.

**Figure 1.**
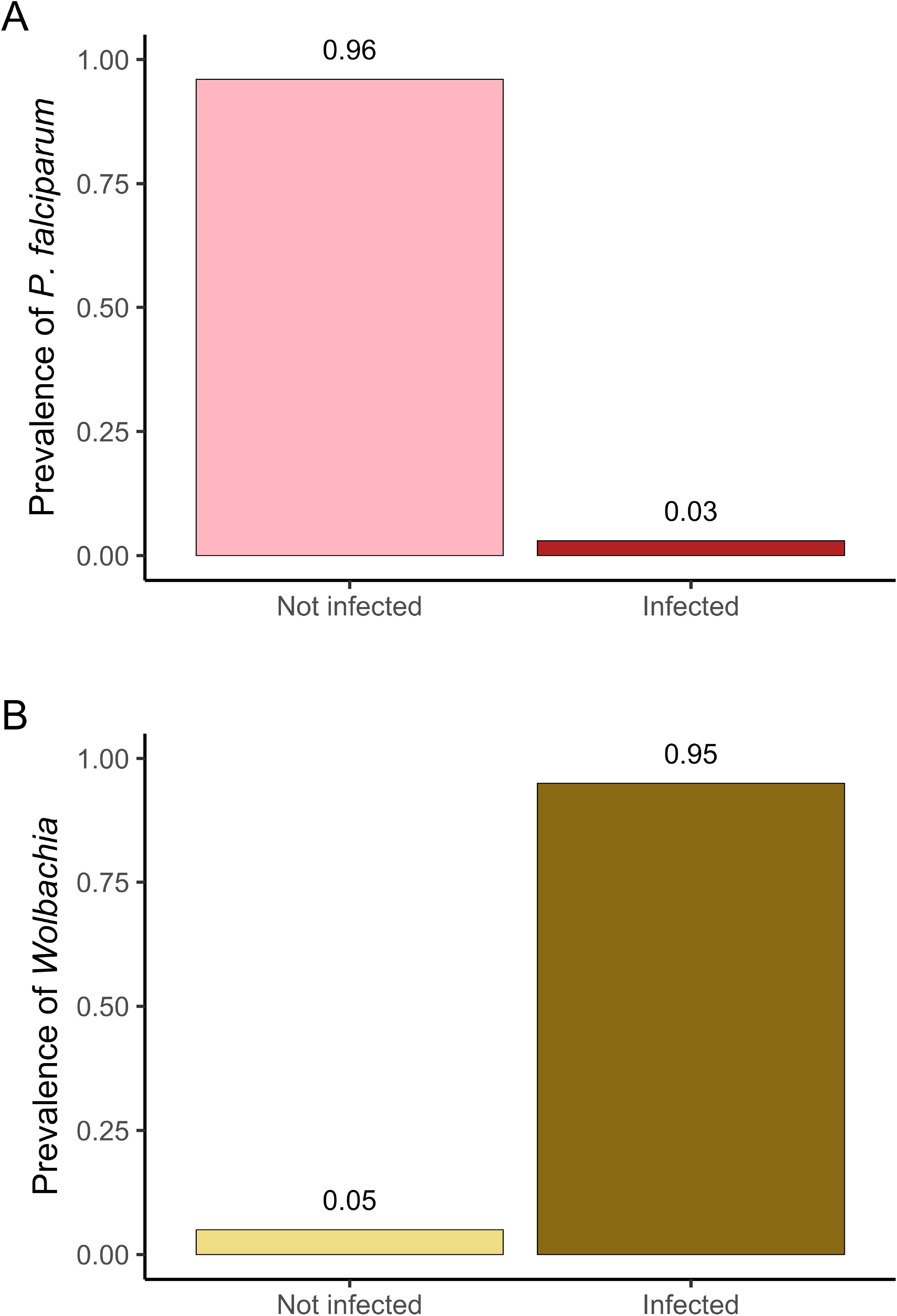
Prevalence of *A. moucheti* infected by *P. falciparum* (A) and *Wolbachia* (A). Number of *A. moucheti* specimens analyzed: 2042 in (A) and 130 in (B).

To estimate the reliability of our quantifications using the plasmid, we performed a repeatability analysis (Stoffel et al., 2020). We randomly re-analyzed 21 mosquitoes from the 64 infected mosquitoes both organisms in this study. In our mixed effects model, we used the ratio of *Wolbachia* and *P. falciparum* value as explanatory variables, while the sample was used as random variable. The number of parametric bootstraps was 1000 for Gaussian data. The repeatability index (R) values were 0.853 for *Wolbachia* (p<0.001) and 0.701 *P. falciparum* (p<0.001).

### Effect of Wolbachia on P. falciparum infection

Higher *Wolbachia* densities are usually observed in the ovaries of its hosts (Werren et al., 2008). To confirm that *Wolbachia* was present also in other *An. moucheti* tissues/organs, we compared the ratio of *Wolbachia coxA* gene copies to mosquito *GPDG* gene copies in whole mosquitoes (n=93) and in abdomen alone (n=14) (Fig. 2). The ratio was significantly higher in whole mosquitoes (Mann-Whitney, W=1387, p < 0.001). Specifically, the median ratios of *Wolbachia* gene copies to mosquito gene copies were 2.02 (maximum ratio=36.93) in whole mosquitoes and 0.0006 (maximum ratio=9.2) in dissected mosquitoes. When looking at *P. falciparum* infection, we found that only two dissected mosquitoes were infected. Conversely, the median ratio of *P. falciparum Cox1* gene copies to mosquito gene copies in whole mosquitoes was 0.17, with a maximum of 8.62, which was much lower than the ratio for *Wolbachia* gene copies.

**Figure 2.**
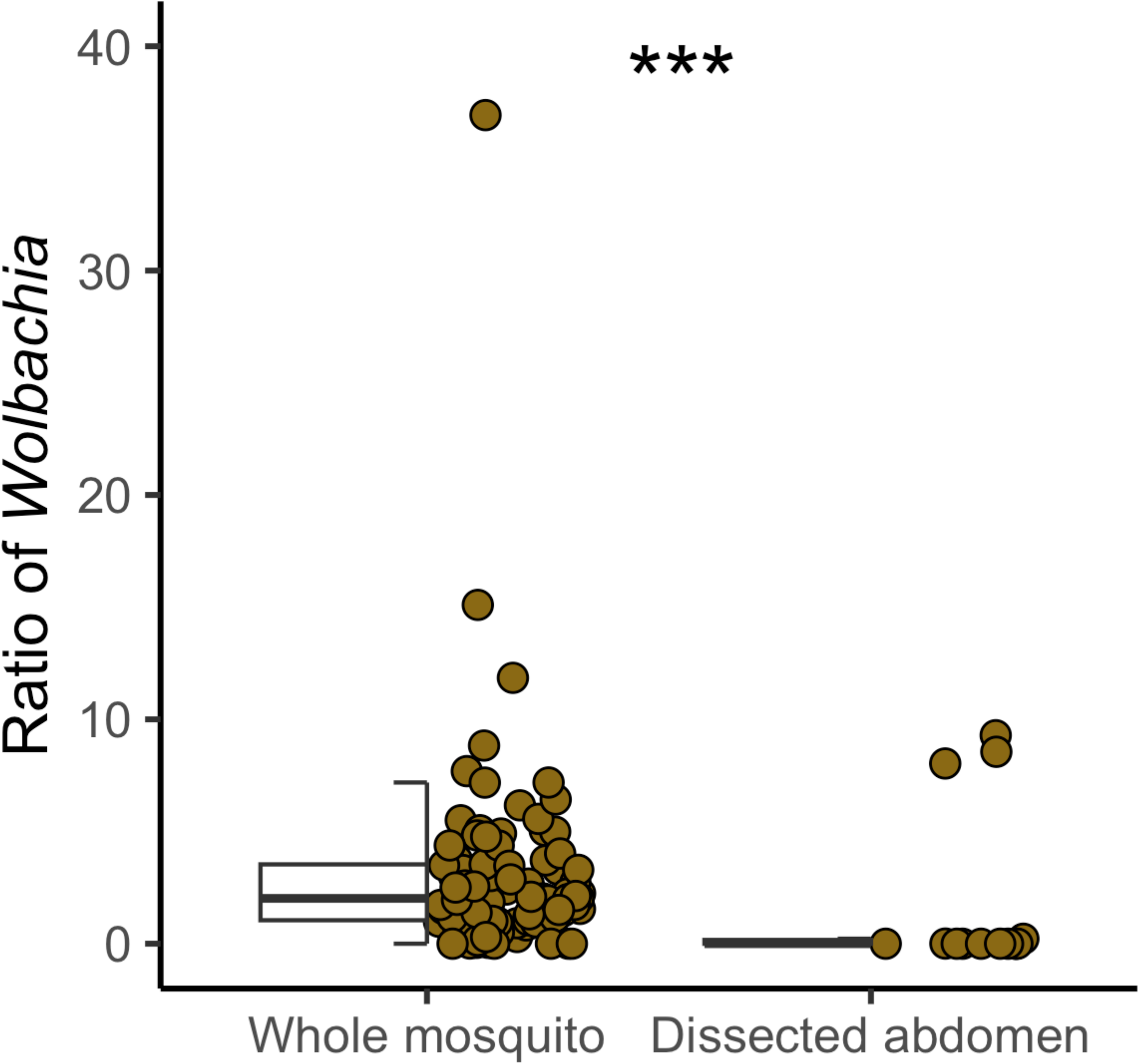
*Wolbachia* gene copies in *A. moucheti* (n=93 whole mosquitoes; n= 14 dissected abdomens). ***p < 0.001 (Mann-Whitney)

Prior studies suggested a potential role of *Wolbachia* in protecting *An. gambiae* from *P. falciparum* infection in natural populations (Gomes et al., 2017; Shaw et al., 2016). We first carried out a qualitative analysis to compare *Wolbachia* density in mosquitoes infected (n=38) and uninfected with *P. falciparum* (n=55) (Fig. 3A). We found that *Wolbachia* density was slightly, but significantly higher in infected (mean gene copies = 4.21) than in uninfected (mean gene copies = 2.23) *An. moucheti* (Mann-Whitney, W=996, p= 0.004). We then asked whether the *Wolbachia*-*P. falciparum* relation was density-dependent. Using the ratio of *Wolbachia* and *P. falciparum* copies relative to their host (*An. moucheti*), we observed a negative, but not significant correlation (Pearson, R=-0.11, p=0.49, Fig 3B). These results suggest that *Wolbachia* does not hinder *P. falciparum* infection or development in this malaria vector, in agreement with its prominent role as major malaria vector in Central Africa (Fontenille & Simard, 2004).

**Figure 3.**
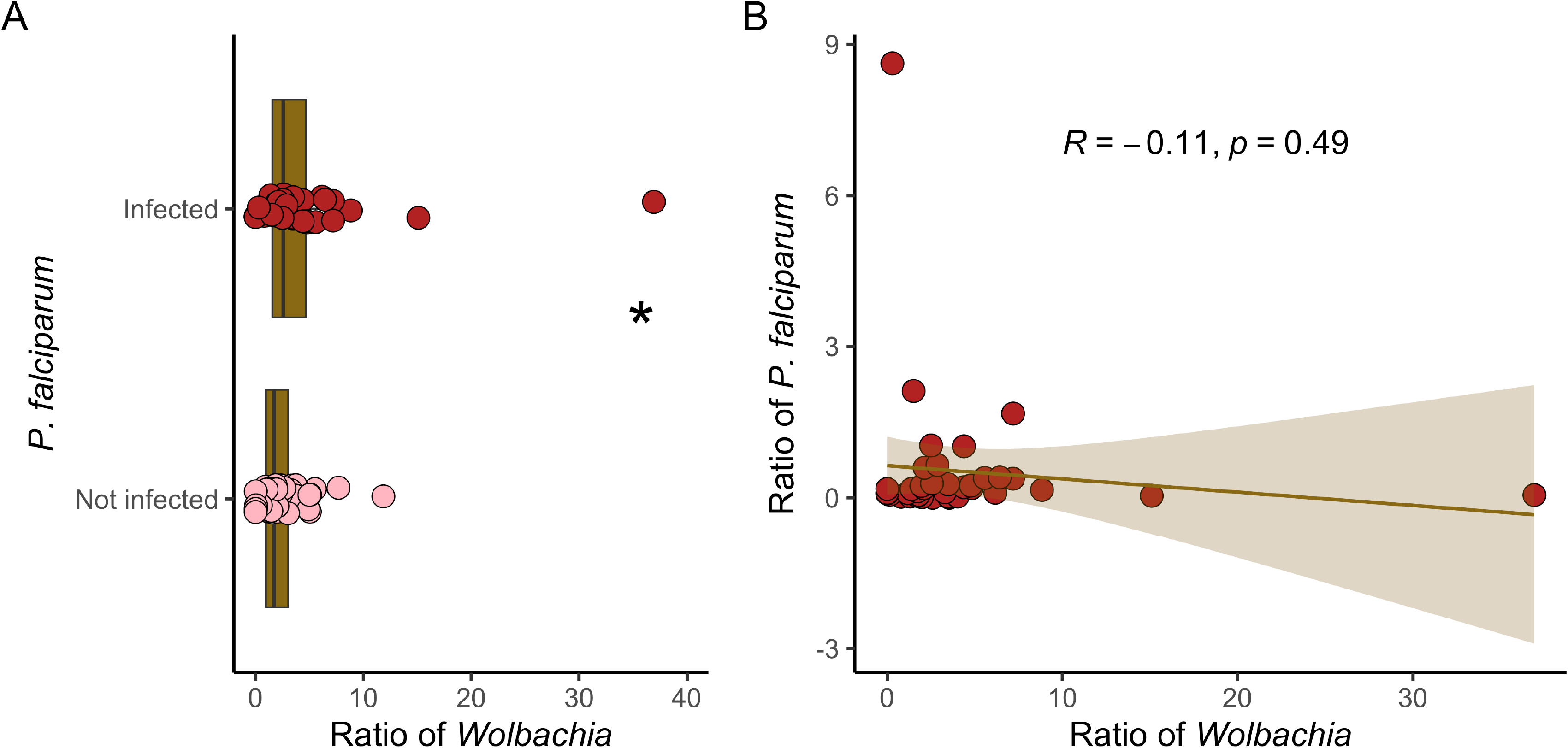
Association of *P. falciparum* and *Wolbachia* in *An. moucheti*. (A) Quantitative analysis of *P. falciparum* (infected vs non-infected) relative to *Wolbachia* gene copies (n=38 *P. falciparum*-infected specimens and n=55 *P. falciparum* non-infected specimens); *p= 0.004 (Mann-Whitney). (B) Correlation (Pearson) between the gene copies of *P. falciparum* and *Wolbachia* in *A. moucheti* (n=113).

## Discussion

*Wolbachia* is a maternally inherited endosymbiont proposed to reduce the malaria burden (Ross et al., 2019). In this study, we evaluated *Wolbachia* protective effect in *An. moucheti*, one of the major malaria vectors in Africa (Christophe Antonio-Nkondjio et al., 2002; Fontenille & Simard, 2004). First, we estimated the number of mosquitoes infected with *Wolbachia* and *P. falciparum* under natural conditions. Second, we quantified *Wolbachia* and *P. falciparum* densities in *An. moucheti* and found that 3% (64/2042) of mosquitoes were infected by *P. falciparum* and 95% (107/113) by *Wolbachia*. We found that *Wolbachia* did not hinder *P. falciparum* infection in *An. moucheti*. Overall, this study allows better understanding *Wolbachia* place as a biological tool for malaria control in *An. moucheti*.

*An. moucheti* presents one of the highest *Wolbachia* prevalence rates in *Anopheles* (Ayala et al., 2019; Jeffries et al., 2018). *Wolbachia* is common across its geographical range (i.e. Cameroon, Gabon, Democratic Republic of Congo), ranging from more than 50% in Cameroon or Democratic Republic of Congo to almost fixed (95%, Fig 1B). Differences in prevalence can be due to geographical heterogeneities, previously observed in other species (Ahmed, Araujo-Jnr, Welch, & Kawahara, 2015), or technical issues. For instance, we used *coxA* instead of 16S for PCR-based detection (Gomes et al., 2017; Jeffries et al., 2018). Nonetheless, our high prevalence is in agreement with results on maternal transmission in Gabon (from 90% to 100%) and with the theoretical prevalence of *Wolbachia* across its hosts (Zug & Hammerstein, 2012). We then quantified *Wolbachia* density in its mosquito host. We observed a median density of 2.02 gene copies of *Wolbachia* in *An. moucheti*. This number is similar to what observed in *Culex pipiens*, a mosquito with high infection rate (Berticat, Rousset, Raymond, Berthomieu, & Weill, 2002). Walker et al., also quantified *Wolbachia* density in *An. moucheti* (Walker et al., 2021). Unfortunately, they used a different method (i.e., *Wolbachia* 16S copies/ng DNA), making impossible the comparison between studies. Another study in *An. gambiae* proposed to normalize the host genome level using the S7 rRNA gene and two independent RT-qPCR assays (Gomes et al., 2017), with similar results. We observed that *Wolbachia* densities varied greatly in our sample. The host age and the physiological status can greatly influence its density, as usually observed in other insect species (Duron, Fort, & Weill, 2007; Tortosa et al., 2010; Unckless, Boelio, Herren, & Jaenike, 2009). Our *An. moucheti* mosquitoes were from a natural population, therefore, without control of age or physiological cycle, and this may explain the variation of *Wolbachia* densities we observed. Moreover, as mosquitoes were kept alive for at most 10 days, some of them could have developed eggs, possibly increasing *Wolbachia* density.

In Central Africa, *An. moucheti* plays a key role in malaria transmission (Christophe Antonio-Nkondjio & Simard, 2013). Therefore, our first question was how this mosquito can be a major malaria vector despite this high *Wolbachia* prevalence if this endosymbiont exerts a negative impact on *P. falciparum* development as previously observed in *An. gambiae* (Gomes et al., 2017; Shaw et al., 2016). The *P. falciparum* infection rate (3%) in our *An. moucheti* population was relatively higher compared with other studies (Christophe Antonio-Nkondjio et al., 2002). This could be explained by our experimental protocol: the probability of developing the infection would increase in mosquitoes that live longer (Dawes, Churcher, Zhuang, Sinden, & Basáñez, 2009). Moreover, we quantified *P. falciparum* density in mosquitoes, and confirmed that in natural conditions, it is very low (Medley et al., 1993). When we studied *Wolbachia* effect on *P. falciparum* development, we first quantitively analyzed *P. falciparum* and *Wolbachia* infection (Fig 3A). Our data suggests that *Wolbachia* density does not affect malaria transmission by this mosquito. Rather, our results revealed that *Wolbachia* density was higher in *P. falciparum-*infected mosquitoes. Only two *P. falciparum-*positive mosquitoes were not infected with *Wolbachia*. Although this represents 33% of the 6 non-*Wolbachia* infected mosquitoes (Table S1), it is much higher than the expected probability (0.15% expected vs 33% observed) to find a non-infected *Wolbachia* mosquito (5%) with *Plasmodium* parasites (3%) (expected probability = 0.05 × 0.03 =0.0015). Unfortunately, due to the low number of uninfected *Wolbachia* specimens, we cannot exclude a random effect. Moreover, the *Plasmodium* density was similar between the non-infected and infected *Wolbachia* mosquitoes (Table S1). In *An. gambiae*, two independent studies revealed a blocking role of *Wolbachia* in *P. falciparum* transmission in natural conditions (Gomes et al., 2017; Shaw et al., 2016). *Wolbachia* protective phenotype is very common when an exogenous *Wolbachia* strain invades a new host (Ross et al., 2019). It has been observed in *An. stephensi* and *An. gambiae*, where transfection of the *Wolbachia* strain (*wAlbB*) drastically reduced the ability to transmit the malaria parasites (Bian et al., 2013; Grant L. Hughes et al., 2011). For instance, this phenotype has boosted the use of the *Wolbachia* strain *w*Mel, coming from the fruit fly, in *Ae. aegypti* (Schmidt et al., 2017; Utarini et al., 2021). On the other hand, in longtime established and high prevalence *Wolbachia*-host associations, it is more unusual, and when it occurs, the bacterium prevalence is low (Zele et al., 2014). Besides hindering *Plasmodium* infection, *Wolbachia* might also affect its development. In our study the *Wolbachia*-*P. falciparum* density correlation was not significant (Fig 3B). Conversely, Gomes et al., showed a significant association between *Wolbachia*, even at low density, with the malaria parasite in *An. gambiae* and *An. coluzzii* (Gomes et al., 2017). This was particularly true in the artificially infected *An. coluzzii* colony with the development of sporozoites (Gomes et al., 2017). We could not differentiate between sporozoite and oocyst infections. In our sampling, only one dissected mosquito (only sporozoites, Table S1) was positive for both *Wolbachia* and *Plasmodium* infection. Altogether, our results indicate that in *An. moucheti, Wolbachia* does not have a protective role against *P. falciparum* infection. These results are in agreement with its main malaria vector role in Central Africa.

Our finding may suggest that natural infections of *Wolbachia* are not useful for malaria control in *An. moucheti*, unlike what proposed by previous studies in *An. gambiae* (Ross & Hoffmann, 2021). Multiple factors could explain these discrepancies. First, while *An. gambiae* has been rarely found infected by *Wolbachia* with prevalence rates higher than 20%, *An. moucheti* exhibits one of the highest prevalence rates among the studied *Anopheles* species(Ayala et al., 2019; Jeffries et al., 2018). Therefore, we could hypothesize that *An. moucheti* immune system has been adapted to *Wolbachia* presence for long time. Actually, the bacterium may increase the transmission in a long-term association as it has been observed in the avian malaria system, where *Wolbachia*-infected *Culex pipiens* mosquitoes transmit better than non-infected (G. L. Hughes, Rivero, & Rasgon, 2014; Zele et al., 2014). Moreover, high *Wolbachia* prevalence is rather associated with cytoplasmic incompatibility than with protection against vector-borne pathogens (Werren et al., 2008). Unfortunately, crossing experiments cannot be performed in *An. moucheti* because of the absence of laboratory colonies. The results of experiments using other *Anopheles* species with established laboratory colonies showed that colonies were infected with slightly different *Wolbachia* strains (Ayala et al., 2019; Jeffries et al., 2018; Walker et al., 2021), which could indicate an unidirectional CI (i.e., *Wolbachia* infected male mated with *Wolbachia* uninfected female), as evidenced in other systems (Werren et al., 2008). On the other hand, in hosts with low *Wolbachia* prevalence, this endosymbiont can be associated more with a protective phenotype(Oliver, Degnan, Burke, & Moran, 2010). This is the case in *An. gambiae* (Gomes et al., 2017). Despite the protective phenotype of the transinfected or naturally occurring *Wolbachia* strains against *Plasmodium* infection, currently, no program has been implemented in which *Wolbachia*-infected *Anopheles* are used to reduce the malaria burden. The main challenge is to find a Wolbachia strain that can invade and exhibit a protective phenotype against malaria infection. The highly prevalent and *Anopheles*-adapted *w*Anmo *Wolbachia* strain could represent an opportunity to carry out transinfections in *Anopheles gambiae*, for instance. More studies on *An. moucheti* and *Wolbachia* are needed to determine the potential value of *Wolbachia* as a biological tool for malaria control.

## Conclusions

The use of *Wolbachia* as vector control strategy is a reality. In the last decade, hundreds of thousands of *Wolbachia*-infected *Ae. Aegypti* specimens have been released in many different countries. As result, several dengue epidemics have been reduced and the vector population size controlled (Utarini et al., 2021). In *Anopheles*, many studies have been conducted to determine *Wolbachia* place as a vector control strategy. Here, we characterized *Wolbachia* role in malaria transmission by *An. moucheti*, one of the major malaria vectors in Africa. Although, *Wolbachia* presence and density were not associated with protection against *P. falciparum* in *An. moucheti*, this strategy could be proposed for other malaria vectors, such as *An. gambiae*.

## Acknowledgements

We would like to thanks the volunteers and authorities from Ndangueng, Cameroon, for their assistance and compliance during the field investigations. We also thank the IRD Representation in Yaoundé for their logistic assistance during this study. This work was funded by ANR, France (ANR-18-CE35-0002-01–WILDING), grant awarded to DA. This project has been funded in whole or in part with Federal funds from the National Institute of Allergy and Infectious Diseases, National Institutes of Health, Department of Health and Human Services, under Grant Number U19AI110818 to the Broad Institute (DEN).

## Supplemental Material

Table S1. List of the *An. moucheti* mosquitoes used in this study.

Text S1. Plasmid sequence.

