## Supplementary material for "No *Wolbachia* protection against *Plasmodium falciparum* infection in the major malaria mosquito *Anopheles moucheti*": Text S1

**Text S1.** Plasmid construction

AAGTTGTTTCCGGACGTTTGATCGTACGGCGTTACATTTGTTTTTAGAGTGCTATACTTACACTGTGTTCAAACAGCGCAGTTCCATCTATCACCATTAATCTATCCGACGTTACATCAGGAATGTTATTGCTAACACTACCGGTTTTAACTGGAGGAGTATTAATGTTATTATCAGACTTACATTTTAATACTTTATTTTTTGACCCAACATTTGCAGGAGATCCAATATGGTGCTATAACTATGCTGCTAACTGATCGCAATATTGGTACTTCCTTTTTTGATCCTGCTGGTGGCGGTGATCCTGTGTTATTTCAACACCTGTTTTGGTTTTTTGGTCATCCAGAAGTTTACATA
